# Rapid Computation of High-Level Visual Surprise

**DOI:** 10.1101/2025.06.20.660166

**Authors:** David Richter, Paula Pena, María Ruz

## Abstract

Predictive processing theories propose that the brain continuously generates expectations about incoming sensory information. Discrepancies between these predictions and actual inputs, sensory prediction errors, guide perceptual inference. A fundamental yet largely unresolved question is which stimulus features the brain predicts, and therefore, what kind of surprise drives neural responses. Here, we investigated this question using EEG and computational modelling based on deep neural networks (DNNs). Participants viewed object images whose identity was probabilistically predicted by preceding cues. We then quantified trial-by-trial surprise at both low-level (early DNN layers) and high-level (late DNN layers) visual feature representations. Results showed that stimulus-evoked responses around 200ms post-stimulus onset over parieto-occipital electrodes were increased by high-level, but not by low-level visual surprise. These findings demonstrate that high-level visual predictions are rapidly integrated into perceptual inference, suggesting that the brain’s predictive machinery is finely tuned to utilize expectations abstracted away from low-level sensory details to facilitate perception.

## Introduction

Perception combines sensory input with prior knowledge and the expectations that we derive from this prior knowledge [1–3]. Accordingly, predictive processing theories postulate that the brain continuously generates predictions about such inputs, resulting in prediction errors, the discrepancies between expected and actual inputs [4–6]. A critical question in understanding predictive processing is: What stimulus features are predicted by the brain? And consequently, what kind of mismatches do sensory prediction errors reflect? One hypothesis, based on classical models of hierarchical predictive coding, is that prediction errors are locally computed in terms of features represented in each visual cortical area. As such, low-level surprise (e.g., unexpected edges and contrasts) is computed in early visual areas, whereas high-level visual surprise (e.g. object features) is computed in higher visual areas, such as in the fusiform gyrus. An alternative hypothesis is that sensory prediction errors largely reflect high-level surprise, relayed from higher-order to early visual areas. The rationale is that surprise is computed in downstream areas and the resulting prediction error, or a scalar quantity reflecting the magnitude of the surprise, modulates neural responses in earlier areas via feedback.

In support of the second account, recent functional Magnetic Resonance (fMRI) evidence suggests that prediction errors across ventral visual areas, including in primary visual cortex (V1), increase with high-level visual but not low-level surprise [7]. Similarly, neural firing rates in macaque V1 scale with high-level image predictability [8]. Combined, these studies suggests that predictions may largely operate at higher levels of abstraction, and consequently that prediction errors may be computed in higher visual areas and subsequently fed back to earlier visual areas. Complementary results demonstrate representations of high-level features in early levels of the face processing hierarchy during prediction [9]. In sum, high-level predictions may influence neural activity across the visual hierarchy.

However, little is known about the timing of these predictive processes, as key studies employed fMRI [7,9]. Yet, understanding when high-level surprise modulates sensory or even post-perceptual processes is crucial, as it provides critical insights into the neurocomputational principles and role of predictions in shaping perception. Early modulations would imply that high-level expectations rapidly influence sensory processing, integrating high-level priors with key perceptual mechanisms. In contrast, late modulations may indicate that such priors only modulate later stages related to updating priors or post-perceptual processes, such as decision-making. If prediction indeed plays a critical role in perception, as proposed by predictive processing theories [4–6], we would expect that high-level surprise modulates key sensory responses, including early to intermediate stages of object perception.

To address the temporal dynamics underlying perceptual prediction, we exposed participants to object images that were expected or unexpected based on preceding cues, while recording electroencephalography (EEG). Using a visual deep neural network (DNN) we quantified low- and high-level visual surprise per trial by computing the representational distances between a seen unexpected stimulus and the stimulus that was expected given the cue. Results showed that high-level visual surprise explained modulations of the stimulus evoked neural activity over (parieto-)occipital electrodes starting ~190 ms post stimulus onset. Specifically, responses were upregulated the more surprising a seen stimulus was in terms of its high-level visual features, but not by low-level visual surprise, suggesting that high-level predictions play an integral role during visual inference.

## Results

To investigate how predictions shape the temporal dynamics of perception, we exposed participants to cue-stimulus pairs, where the letter cue probabilistically predicted the identity of the object image. A trial is depicted in Figure 1A. Each participant was presented with a different set of letter cue and object image combinations. Eight object images were selected per participant from a set of 233 images ensuring variation in both low-level and high-level visual features (see *Stimuli and Experimental Paradigm* for details and S1 Figure an example stimulus set). Each of those eight images were paired with one letter cue (eight randomly selected consonants). As illustrated in Figure 1B, images were seven times more likely to follow their associated cue than any other image. Each object image served equally often as expected and as unexpected image. Moreover, all images appeared equally often as unexpected stimuli, thus participants were equally familiar with all stimuli. They were not informed about the underlying regularities but acquired them by incidental statistical learning. Participants were tasked to respond as fast and accurately as possible to the object images, classifying them as animate or inanimate. Additionally, no-go trials were added, consisting of vowel cues, which required participants to withhold a response. These no-go trials were added to encourage learning of the underlying statistical regularities by making both letter cues and object stimuli task relevant.

**Figure 1.**
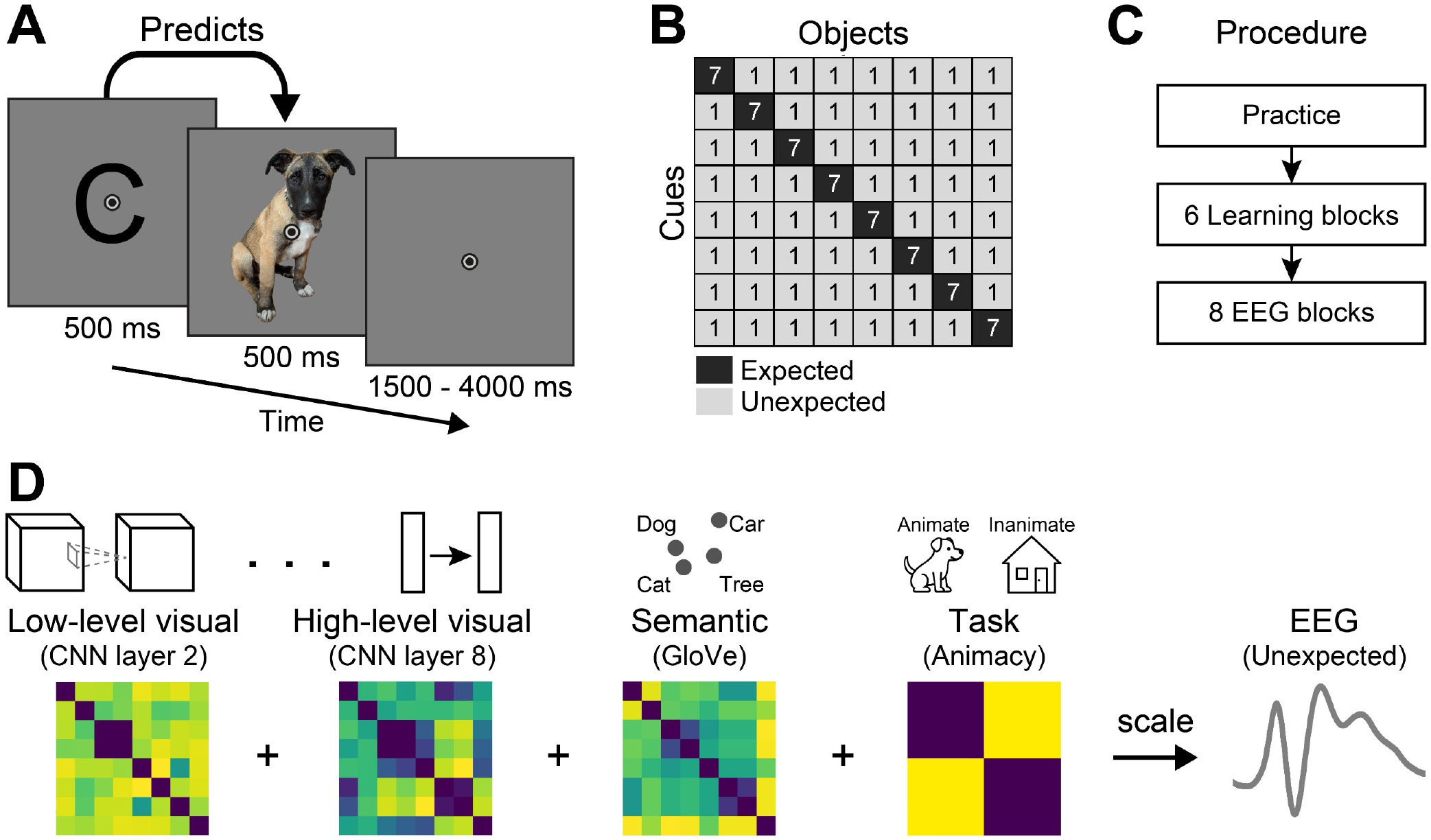
Paradigm and Analysis Rationale. **A)** A single trial, depicting the letter cue (500ms) which probabilistically predicted the object stimulus (500ms). Participants classified images as depicting animate or inanimate entities. **B)** Transitional probability matrix determining the association between letter cues and object images. Numbers indicate trials per block. **C)** Experiment procedure, consisting of practice, 6 learning and 8 main EEG blocks. Associations during learning blocks were strengthened compared to EEG blocks to encourage prediction (for details see: *Materials and Methods: Experimental Procedure*). **D)** Analysis rationale, illustrating the extraction of representational dissimilarity matrices for the eight object images seen by one example participant from an early and late layer of the visual DNN, word embedding, and animacy model. EEG data from unexpected image trials was regressed per timepoint onto the RDMs. The obtained beta coefficients indicate how the EEG signal was scaled by low-level visual (layer 2) and high-level visual (layer 8) surprise, as well as semantic (word level; GloVe) and task (animacy) based surprise.

### Expectations Facilitate Behavioral Responses

Before analyzing the EEG data, we first established whether participants learned and used the statistical regularities underlying the cue-stimulus pairs. Participants behavioral responses were facilitated by expectations both in terms of response speed (Figure 2A; *F*_(2,74)_ = 13.92, *p* < 0.001, 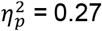) and accuracy (Figure 2B; *F*_(1.6,59.8)_ = 8.62, *p* = 0.001, 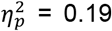). Specifically, the average reaction times (RTs) were faster for expected stimuli (511 ms) compared to unexpected ones, irrespective of whether that stimulus required the same response (516 ms; *t*_(37)_ = 2.08, *p* = 0.045, *d*_*z*_ = 0.05) or a different response (526 ms; *t*_(37)_ = 5.04, *p* < 0.001, *d*_*z*_ = 0.16) as the expected stimulus. Participants were also slower to respond to unexpected stimuli that required a different response (as the expected stimulus) compared to unexpected inputs that required the same response (*t*_(37)_ = 3.07, *p* = 0.008, *d*_*z*_ = 0.11). Combined, these results suggest a response preparation and a perceptual facilitation due to valid prediction. Response accuracy was higher for expected (96.4%) compared to unexpected stimuli requiring a different response (95.1%; *t*_(37)_ = 3.41, *p* = 0.005, *d*_*z*_ = 0.36). No significant accuracy differences were found between unexpected stimuli requiring the same responses (96.1%) as expected inputs and expected stimuli (*t*_(37)_ = 1.14, *p* = 0.261, *d*_*z*_ = 0.08). In sum, behavioral data demonstrate that participants learned and used the underlying statistical regularities to facilitate responses to predictable stimuli.

**Figure 2.**
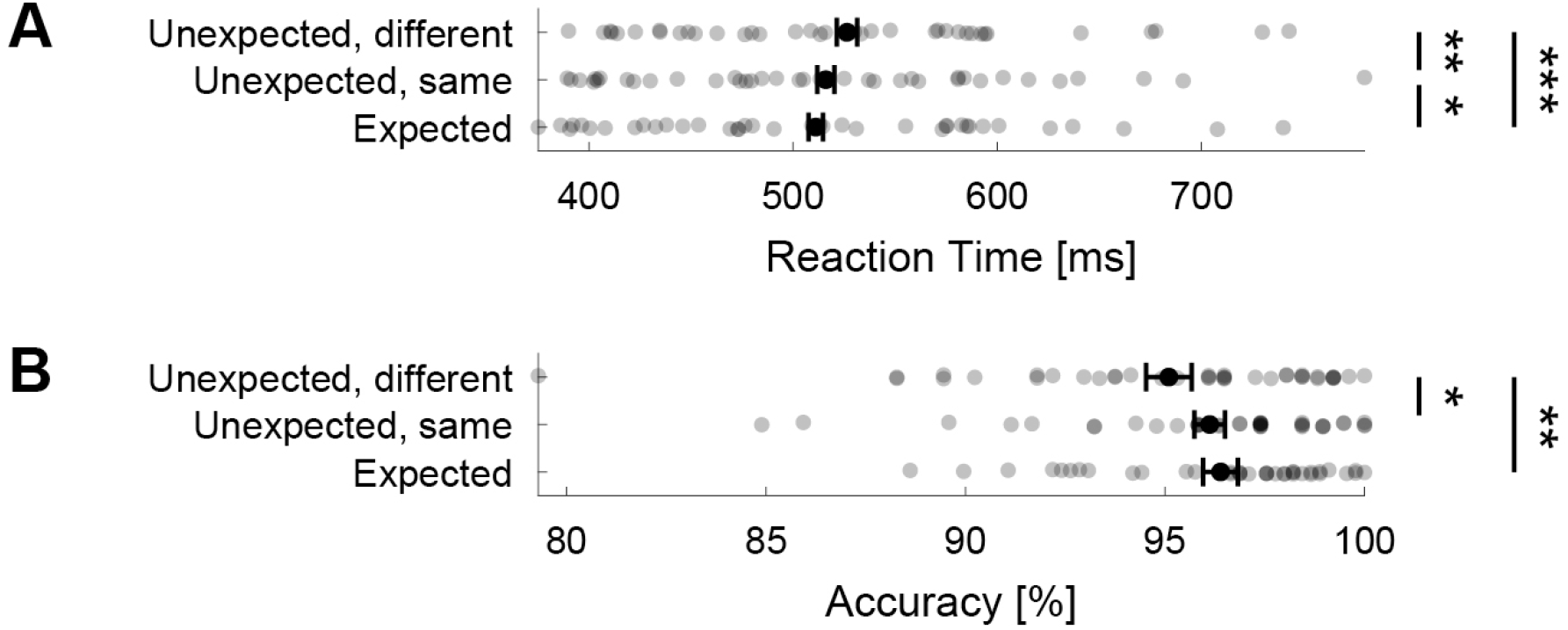
Expectations facilitate behavioral responses. **A)** RTs were faster to expected compared to unexpected stimuli. *Unexpected same* denotes unexpected stimuli that required that same button press as the expected stimuli (i.e., same animacy category). *Unexpected different* are stimuli that required the different button press compared to the expected stimulus. **B)** Response accuracies were higher to expected compared to unexpected stimuli of a different animacy category. Error bars denote within-subject 95% confidence intervals. P values are FWER corrected. *** *p* < 0.001, ** *p* < 0.01, * *p* < 0.05.

### Visual DNNs Explain Visually Evoked Responses

Next, we established whether the utilized visual DNN models explain visually evoked responses in the EEG. We considered this step essential, as the model used to derive low-level and high-level visual surprise also ought to explain generic visual responses. To this end, we conducted a representational similarity analysis (RSA [10]). In brief, for each participant we formed the neural representational dissimilarity matrix (RDM) by computing the pair-wise representational distance (1 – correlation) between stimuli across all EEG channels per timepoint, irrespective of expectation status. Next, we derived multiple model RDMs by computing the representational distances between the stimuli from the layers of interest of the visual DNN (layer 2 and 8), as well as a word embedding based semantic model (GloVe), and a task model using animacy category. As a control model we also included an untrained (randomly initialized) version of the DNN layer 8. Finally, we computed the correlation (Kendall’s Tau) between these model RDMs and the neural RDM per timepoint. Thus, the resulting correlation indexes the shared representational geometry between the model RDMs and the EEG signal.

RSA results, depicted in Figure 3A, showed that all models, except for the untrained layer 8 control, explained the neural response. Critically for our primary analyses, both the low-level (layer 2; 74– 640 ms, *t*_max_ = 5.94, *p*_cluster_ < 0.001) and the high-level visual model (layer 8; 98–594 ms, *t*_max_ = 7.70, *p*_cluster_ < 0.001) explained the visually evoked response in a broad time-window following stimulus onset. Numerically the estimated onset time for the low-level model (79 ms) was earlier than for the high-level model (107 ms), however this difference in timing did not achieve statistical significance (*p* = 0.077). The semantic, non-visual model (GloVe; 86–570 ms, *t*_max_ = 7.23, *p*_cluster_ < 0.001) and the task model (animacy category; 102–645 ms, *t*_max_ = 5.38, *p*_cluster_ < 0.001) also reliably explained the EEG responses in a wide time window. In sum, these results confirm that the utilized models properly explain the visually evoked responses as measured by EEG in appropriate time-windows.

**Figure 3.**
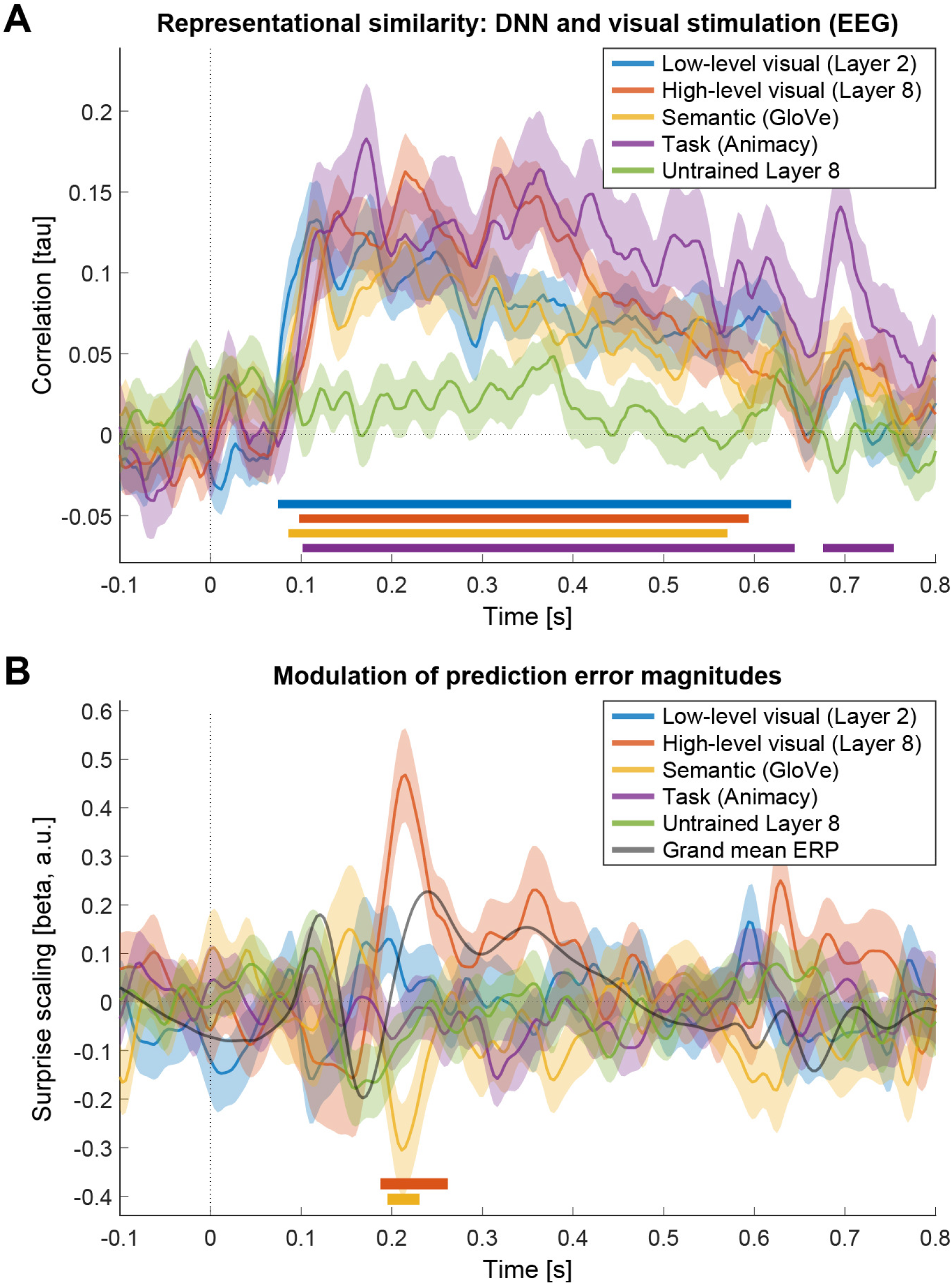
**A)** Representational similarity analysis shows that the utilized models explained visually evoked EEG responses. All models of interest, low-level visual (layer 2), high-level visual (layer 8), a word-based model (GloVe), and task model (animacy category) explained evoked responses shortly after stimulus onset. RSA was computed across all electrodes. **B)** Visually evoked EEG responses scaled with high-level surprise approximately 200 ms post stimulus onset. In contrast to A, the visually evoked response, only high-level visual and semantic surprise modulated the magnitude event related potentials. Because our focus was on visually evoked ERPs, the depicted results were averaged across all (parieto-)occipital electrodes (Oz, O1, O2, POz, PO7, PO8, PO3, PO4). For temporal reference, the grey line indicates the grand mean ERP computed over the same parieto-occipital electrodes. Colored bars above the abscissa denote statistically significant clusters (*p*_cluster_ < 0.05).

### High-Level Visual Surprise Modulates Early Visually Evoked Responses

Having established that our DNN models of interest successfully explained evoked responses, we next focused on how sensory prediction errors are modulated by surprise. Specifically, we assessed whether the amplitude of visual ERPs elicited by unexpected stimuli over parieto-occipital electrodes was modulated by different levels of surprise. In brief, surprise was indexed using the representational distance of a seen unexpected stimulus from the stimulus that was expected on a trial given the preceding cue. Thus, we quantified surprise on a trial-by-trial level contingent on the expectation, induced by the statistical regularities (Figure 1B) and the unexpected seen image. High-level visual surprise was derived from a late layer of the DNN (layer 8) and low-level surprise from an early layer (layer 2) based on previous work [7]. A non-visual task-surprise metric was derived from the animacy category and a non-visual semantic surprise model using word embedding (GloVe), again using the representational distance between the seen unexpected stimulus compared to the expected input. Finally, untrained (randomly initialized) layer 8 distances were included as a control model. Surprise, as indexed by these 5 models, was then regressed onto the EEG response using multiple linear regression (Figure 1D). Therefore, the resulting betas indicate how the evoked responses are up or down regulated as a function of surprise. Because our focus was on visual prediction error computations, we analyzed all occipital and parieto-occipital electrodes (Oz, O1, O2, POz, PO7, PO8, PO3, PO4).

Figure 3B shows that surprise modulated neural responses over parieto-occipital electrodes approximately 200 ms post-stimulus onset (onset time = 191 ms), corresponding to the visual P2 potential. Specifically, we found a statistically significant cluster reflecting an increase of the evoked response by high-level visual surprise (layer 8: 188– 262 ms, *t*_max_ = 5.82, *p*_cluster_ = 0.001). One additional statistically significant cluster was observed, suggesting a downregulation of evoked responses by semantic (word based) surprise (GloVe: 195–230 ms, *t*_max_ = −4.06, *p*_cluster_ = 0.046; onset time = 192 ms). Interestingly, no other metric, including low-level visual surprise (layer 2), attained statistical significance, suggesting that early to intermediate visual processing is predominantly modulated by high-level (visual) surprise. Notably, we did not observe any significant modulations of late evoked responses (>300 ms post stimulus onset) by any surprise metric over parieto-occipital sensors. To address potential concerns about correlated regressors, we repeated the analysis using Shapley value regression (S2A Figure) [11]. Additionally, to ensure that our results were not contingent on the a priori selected DNN layers, we performed the analysis using adjacent DNN layers (S2B Figure). Both control analyses replicated the results reported above, suggesting that our results are robust to the specific analysis methods and DNN layer selection.

To allow for interindividual variability in peak ERP responses, we performed an additional analysis with time-windows specified for each participant. In brief, for each participant and visually evoked ERP, we defined a 20 ms window around the maximal individual deflection (negative or positive depending on the ERP) within broad time-windows commonly reported in the literature for each ERP; P1: 80-130 ms; N1/N170: 130-190 ms; P2: 180-250 ms; N2: 200-300 ms, P3: 300-600 ms post-stimulus onset [12–25]. Within the 20 ms window around the individual peaks we averaged the beta estimates for each participant and ERP, and finally computed the mean across participants.

Results of these time-window ERP analyses are depicted in Figure 4. First, in agreement with the time-resolved results, P2 amplitudes were significantly increased by high-level visual surprise (layer 8; *t*_(37)_ = 5.46, *p* < 0.001, *d*_*z*_ = 0.89). Additionally, P2 amplitudes were significantly more upregulated by high-level visual surprise compared to all other surprise metrics (all paired t-tests *p* < 0.01; see S1 Table for details). We did not find any additional modulation by other surprise metrics during any ERP time window (all one-sample t-tests *p*_corrected_ > 0.05; see S2 Table for details). Critically, this included no modulation of neural responses by low-level surprise (layer 2) for any tested ERP – indeed for the N1, P2, and P3 there was evidence for the absence of a modulation of ERP amplitudes by low-level surprise (BF_10_ < ⅓). Moreover, we did not see any modulation of the earliest visual ERPs (P1, N1) by any model of interest. Only the control model (untrained layer 8) resulted in slightly reduced N1 amplitudes compared to the layer 2 and GloVe models, however given the nature of this model and the lack of a reliable difference compared to zero this likely represents a false positive due to random fluctuations. Finally, we found an additional small modulation of N2 amplitudes by layer 8 compared to the Glove and untrained layer 8 models, however again in the absence of a reliable contrast against zero, these results may not reflect a reliable effect. In sum, results of the participant specific time-window analysis across early to intermediate visually evoked ERPs confirmed our key results (Figure 3B) showing a reliable upregulation event related responses by high-level visual surprise approximately 200 ms after stimulus, evident in the modulation of P2 amplitudes.

**Figure 4.**
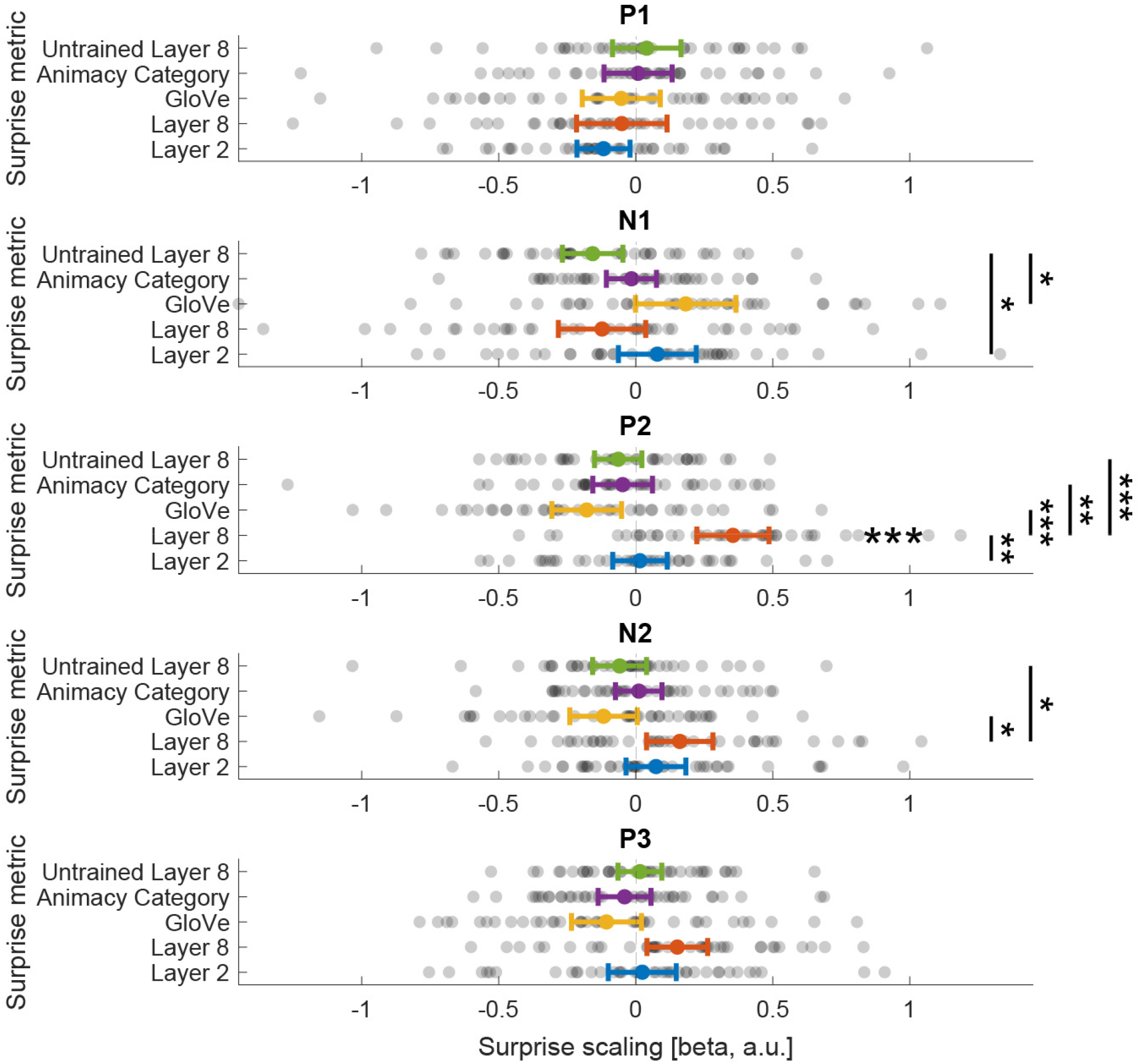
Participant specific time window ERP analysis. For each participant we defined individual ERP peaks within commonly reported time windows. Surprise metrics: Layer 2 (Low-level visual), Layer 8 (High-level visual), GloVe (Semantic), Animacy category (Task), Untrained Layer 8 (Randomly initialized control model). P values are FDR corrected. *** *p* < 0.001, ** *p* < 0.01, * *p* < 0.05.

## Discussion

Predictive processing theories [4–6] hold that prior knowledge, and the predictions derived from this prior knowledge, are key components of cortical processing, including perception. However, what the brain predicts during vision remains largely unknown. One possibility is that the brain predicts across levels of abstraction – that is, in terms of visual features represented across the various visual cortical areas. On this account, V1 for example would predict and hence represent surprise in terms of low-level visual features (e.g., orientation, spatial frequency) with increasingly complex predictions in later visual cortical hierarchy. Alternatively, surprise may be computed at a higher level of abstraction (e.g. object parts and textures), possibly in higher-order visual areas, and the magnitude of this surprise is relayed to early visual cortex. This latter account would thus abstract beyond low-level sensory regularities, instead favoring prediction about high-level features. Arbitrating between these accounts is essential as it provides critical details of the neural mechanisms underlying visual inference. Here we provide new insights into the neurocomputational principles underlying perception by investigating the temporal dynamics underlying perceptual prediction using EEG and surprise quantified using a DNN.

### Rapid Integration of High-Level Predictions during Visual Perception

Following incidental visual statistical learning of object image predictions, we found that high-level visual surprise increases evoked neural responses approximately 190–260 ms after stimulus onset over occipital and parieto-occipital sensors. Our results suggest that cortical visual surprise is largely determined by predictions and prediction error computations at higher levels of visual feature abstraction. Thus, the brain seems to be particularly attuned to discrepancies from expectations in terms of high-level visual features.

The timing, starting ~190 ms post-stimulus onset, coincides with the onset of the visual P2 ERP. Modulations of the P2 have been linked to perceptual feedback [17,18,23,26,27] suggesting that the upregulation of the P2 by surprise may involve a rapid integration of high-level predictions into early feedback mechanisms as vision unfolds. While the limitations of EEG prevent precise inference of the underlying cortical sources, previous studies have reported modulations of V1 responses by high-level predictions [7,8,28]. A plausible mechanism accounting for the combined observations may involve direct feedback connections from higher-order visual areas, such as fusiform cortex, to earlier visual cortical areas [29]. Category-selective fusiform activity for instance feeds back to V1 within approximately 150–200 ms [30,31], thus matching the timing of the modulations by surprise seen here. Precisely these later visual areas contain neurons tuned to high-level visual features, making them likely nodes for computing and relaying surprise at this level of abstraction. Thus, it seems plausible that feedback projections to early visual cortex swiftly modulate neuronal activity on the basis of high-level visual predictions, or surprise signals, originating from higher-order visual areas, and possibly extra-visual regions such as the hippocampus [32–34].

### Predictive Processing and High-Level Prediction

The mechanism described above aligns well with hierarchical predictive coding theories [5,6,35,36], where predictions are compared with sensory data, thereby generating prediction errors. In this context, our results suggest that visual prediction errors largely scale with higher-level surprise during perceptual inference. In line with this interpretation, work using fMRI showed that high-level visual surprise increases neural activity throughout the visual system [7]. Our EEG data complement these results by demonstrating that these effects occur relatively early during the visual response. Together, these findings support a model where high-level predictions rapidly influence visual processing across visual cortex, including during early to intermediate stages of visual inference. This integration of spatial and temporal data strengthens predictive coding’s assertion that perception involves rapid interactions between predictions and sensory inputs across hierarchical levels of processing. The integration of high-level predictions may allow for an efficient updating of internal models as perception unfolds, thereby facilitating the interpretation of sensory inputs.

However, the lack of neural modulations by low-level visual surprise, in both the prior fMRI study [7] and the current EEG data, is notable. First, it is important to emphasize that stimuli were predictable in terms of low-level features, thereby in principle allowing for precise low-level predictions. Moreover, the utilized low-level visual feature model well explained visual responses independent of surprise. Therefore, the failure of low-level surprise to scale prediction error magnitudes is unlikely to stem from a general shortcoming of the DNN model or a lack of low-level predictability. Instead, our results suggest that the brain may abstract away from low-level regularities. This conclusion integrates well with previous findings, showing that visual neural responses are modulated by high-level predictions [7–9,28]. Combined with our present study, a range of different recording methods, distinct populations, and different kinds of predictions have been studied, yet all converge on a similar conclusion – the visual brain predicts at higher levels while abstracting away from low-level visual regularities [7–9,28]. These results pose important constraints on the implementation of predictive processing, particularly for accounts requiring the computation of prediction errors across all levels of visual abstraction. Our results do appear to be compatible with different implementations of predictive processing. As discussed in more detail elsewhere [37], these include (1) the computation of high-level error signals in late visual areas, which are subsequently relayed to early visual cortex, (2) early visual areas as a comparator for high-level features, and (3) dendritic hierarchical predictive coding [38,39].

### Task and Levels of Prediction

An open question is whether the prioritization of high-level visual surprise observed here reflect task demands. Participants reported stimulus animacy, a high-level task, which likely encourages more abstract predictions. While we did show that animacy reliably explained neural variance of the visual evoked response itself, animacy *surprise* did not modulate ERP amplitudes, suggesting that task surprise does not explain the present results. Moreover, similar findings, showing high-level visual prediction, have been reported in mice and macaques, in the absence of an explicit high-level task [8,28]. Combined, these results suggest a general propensity of the visual system to predict at higher levels of abstraction irrespective of task.

Why would the brain predict at this level of abstraction? An emphasis on high-level visual prediction could reflect evolutionary advantages. High-level visual predictions may constitute a particularly well-suited level of abstraction to support the rapid detection and hence response to important events. On the one hand it is a visual representation, thereby rapidly computable within visual cortex, but one that likely signifies behaviorally relevant events. Thus, prediction at this level of abstraction may be an adaptive strategy. Additionally, Heilbron and de Lange [28] recently suggested that high-level prediction may be an effective way for self-supervised learning of visual representations, drawing parallels with recent successes in machine learning [40,41].

That said, it is nonetheless plausible that task goals shaped the level of prediction. Indeed, it seems likely that the brain can utilize predictions at lower levels of abstraction, such as predictions of local orientation and contrast. Moreover, prior work showed significant prediction error signals in early visual areas for stimuli mostly comprised of low-level features (e.g., Gabor patches; [42]). However, also a Gabor patch can be abstracted, even to a linguistic level (e.g. “the vertical Gabor patch”). Thus, arguably showing prediction error like responses for stimuli mostly consisting of low-level features does not necessary imply that predictions were made at the level of low-level visual features. Future research is necessary to explore how visual predictions operate under different tasks demands while indexing surprise across different levels of abstraction, thereby elucidating the flexibility and specificity of predictive processes in the visual brain.

### Dissociation of Semantic and High-Level Visual Prediction

A curious result was that semantic surprise, here defined using a word-embedding model (GloVe; [43]), appeared to have an opposite effect on evoked responses compared to high-level visual surprise. That is, we found a reduction of P2 amplitudes by semantic surprise. We note that these results should be interpreted with caution, as the effect did not attain statistical significance in the individual-peak based ERP analysis but only in the time-resolved analysis. Taken at face value, these results suggest that unexpected inputs of a semantic category similar to the expected stimulus would elicit *larger* prediction errors compared to very dissimilar categories – opposite to what one might intuitively expect. Why could this be the case?

One speculative explanation is that while high-level surprise may drive visual prediction errors, semantic predictions may aid in the differentiation of related categories. For example, while a lion might not be very different, and hence not surprising, in terms of its high-level visual features when you expected to see your Leonberger (giant dog), it is nonetheless a critical semantic distinction, which needs to be rapidly evaluated and separated. Thus, on this account, high-level visual surprise primarily scales sensory prediction errors magnitudes, but simultaneously semantic *proximity* may trigger an increased error response to aid in the separation of related but distinct categories. In contrast, semantically distant unexpected inputs may not require this aid, as they are sufficiently distinct in terms of high-level visual features to allow for reliable identification. That semantic priors appear to be integrated during perceptual inference, and even modulate sensory responses in V1, has previously been suggested [44], thereby giving more credibility to the hypothesis that semantic surprise may directly impinge on visual inference. However, we caution that this interpretation is highly speculative given the present evidence.

### Differentiation from Attention

Surprise attracts attention, and hence, surprising stimuli may use more attentional resources than expected ones [45]. Surprising inputs are also more difficult to categorize compared to expected stimuli. Therefore, we may ask whether the present results can be explained by increased difficulty and attention to unexpected inputs [46]. We believe that this is unlikely. First, key analyses did not depend on contrasting neural responses to expected compared to unexpected inputs, instead relying on quantifying surprise elicited by different unexpected inputs. This approach therefore avoids problems with contrasting expected and unexpected inputs [47]. Moreover, our data showed that a task derived surprise model (animacy category) did not explain P2 amplitudes. Combined, it thus seems unlikely that task difficulty explains the present results.

However, this does not rule out that the modulation of the P2 itself reflects attention. It is plausible that surprising high-level visual features are detected in higher visual areas, and a surprise signal is relayed down the visual hierarchy via feedback. This feedback signal may modulate the visual evoked response directly or instead cause a subsequent increase of attention to the surprising input, which in turn results in the upregulated responses [46]. While this leaves open which precise neurocomputational mechanism the upregulation of the P2 amplitude itself represents (a prediction error, response scaling by surprise, or a subsequent modulation by attention), it does not change the key result that high-level visual surprise fundamentally and rapidly shapes neural responses as perception unfolds. Future work is necessary to further disentangle attention from prediction, as the two factors can be delineated conceptually [48,49] and experimentally [50,51].

### Limitations

We did not find any late (>300 ms post stimulus) modulations of neural signals by any surprise metric. Initially this may appear to contrasts against previous reports in the macaque face processing hierarchy, with higher level areas endowing earlier stages with invariant representations during late processing stages more than 400 ms after stimulus onset [52]. However, EEG lacks the ability to reliably detect some modulations readily recorded by electrophysiology in macaques, which could explain the absence of such late modulations in the present data. Moreover, our analyses were stimulus locked (not response locked), which leaves open the possibility that some response and other post-perceptual modulations were not detected. Therefore, our results should not be taken as evidence against the notion that predictions modulate late visual or post-perceptual processing stages as well.

Repetition suppression, arising from repeated exposure to the same stimulus, is frequently conflated with prediction [47], constituting a potential limitation. Similarly, stimulus familiarity has distinct effects on neural responses [53] and can also be confused with expectation [47]. In the present study, all stimuli were shown equally often and with equal repetition frequency within blocks, thereby precluding an account of our results by simple repetition suppression, adaptation, or stimulus familiarity.

While visual DNNs provide a principled and powerful method to establish visual feature spaces across various levels of abstraction [54,55], their use is not without challenges. First, it is essential to establish that the utilized DNN reliably explains neural responses. Using RSA, we showed that the DNN shares representation geometry with the EEG data. Moreover, previously a convergence of the same DNN and visual cortical representations, as measured by fMRI BOLD, from low-to high-level features has been reported [7]. In other words, early layers of the network best explained early visual cortex representations and late layers higher visual cortex representations. Combined, these factors support a meaningful application of the DNN. Second, the interpretation of DNN feature representations remains difficult. The present model has been shown to represent orientation-like features in early layers (layer 2), while later layers (layer 8) reflect complex texture-like features preferentially encoding object information compared to earlier layers. These results support the interpretation of the DNN layers in terms of meaningful low-level and high-level visual features respectively. For more details on the DNN feature visualization see Richter et al. [7] and for more information on the training dataset and DNN architecture Mehrer et al. [55]. Nonetheless, interpreting the precise features represented by the visual DNN, and hence what is predicted by the brain, remains challenging, and future work is necessary to expand on the present results by using complementary methods to define feature spaces.

## Conclusions

In summary, our results reveal that high-level visual surprise swiftly modulates sensory processing, approximately 190–260 ms post-stimulus onset, as evidenced by effects on the visual P2 ERP. These findings suggest that the brain rapidly incorporates high-level expectations into the recurrent process of perceptual inference. Thereby high-level predictions appear to fundamentally influence how sensory information is processed. The lack of low-level surprise modulating evoked responses further emphasizes the dominant role of high-level features in driving cortical surprise computation. This emphasis on a rapid integration of high-level visual prediction is arguably sensible from an evolutionary perspective, as such features may constitute a particularly meaningful level to aid perceptual inference.

## Materials and Methods

### Participants and data exclusion

Data was acquired at the Mind, Brain and Behavior Research Center (CIMCYC) of the University of Granada. We recruited 40 healthy volunteers through the university’s research participation system. Data from 2 participants were excluded based on subpar behavioral performance (see *Data Exclusion*). This resulted in a final sample of *n* = 38 (31 female; mean age = 21.5 years, SD = 2.9 years).

### Data Exclusion

Participants were excluded if their overall accuracy or reaction times were more than three standard deviations below (for accuracy) or above (for RT) the sample mean, indicating lack of attention or task compliance. Trials contaminated with noise were excluded from EEG data analysis; for details see section *EEG Data Acquisition and Preprocessing*.

### Stimuli and Experimental Paradigm

Participants were presented with pairs of letter cues and full-color images from various categories while EEG data were recorded. On each trial, a letter cue probabilistically predicted the identity of the subsequent image. Specifically, each expected image was seven times more likely to follow its associated letter cue compared to any unexpected image. The same images appeared as both expected and unexpected stimuli, with expectation status determined by the preceding cue. The transitional probability matrix is depicted in Figure 1B.

We utilized full-color images selected from a database of 233 photographs, originally compiled for a previous study [44]. The images represented various categories, including animate entities (e.g., dogs, elephants, human faces) and inanimate objects (e.g., cars, guitars, houses, etc.). Outliers were excluded based on hierarchical clustering of image representations derived from layers 2 and 8 of the DNN model; for details see Richter et al. [7]. From the remaining images, we selected eight images for each participant, including four animate and four inanimate. The selection was optimized to maximize within-layer variance of the layers of interest (layer 2 and layer 8 both contributing equally) and minimize the absolute across-layer correlation of the DNN-derived RDMs, enhancing our ability to detect distinct contributions from different feature levels. Critically, the resulting image sets contained considerable natural variation in both low-level and high-level visual features. For the stimulus set and RDMs of an example participant see S1 Figure, while the full set of RDM visualizations for each participant across all DNN layers is available online: https://doi.org/10.17605/OSF.IO/QSRZ4.

Stimulus presentation and behavioral data collection were performed using MATLAB (version 2020a; The MathWorks, Inc., Natick, Massachusetts, United States) with the Psychophysics Toolbox 3 (Brainard, 1997) on a Windows PC. Images were displayed on an LCD monitor with a resolution of 1920 × 1080 pixels and a refresh rate of 120 Hz, presented over a grey background. Images were displayed centrally, subtending a maximum visual angle of 6°×6°, with actual size depending on the object’s shape. A fixation bull’s-eye (outer circle diameter of 0.5° visual angle) was overlaid at the center of each image to maintain participants’ gaze.

### Experimental Procedure

Each trial began with a letter cue presented for 500 ms, immediately followed by an image displayed for 500 ms, without interstimulus interval between the cue and the image. Participants were instructed to categorize the entity in the image as animate or inanimate as quickly and accurately as possible by pressing one of two buttons. Response mappings were counterbalanced across participants.

To encourage attention to the letter cues and facilitate statistical learning, participants were instructed to withhold their response (no-go trials) when the letter cue was a vowel (a, e, i, o, u). No-go trials were excluded from analysis. Trials were separated by an intertrial interval averaging 2500 ms (range: 1500 – 4000 ms; sampled from a truncated exponential distribution), during which only the fixation bull’s-eye was displayed.

Each experimental block consisted of 128 trials (~7 minutes). The transitional probability matrix (Figure 1B) was presented once per block, resulting in each unexpected cue-image pair being shown exactly once (56 unexpected trials) and each expected pair shown seven times (56 expected trials). The remaining 16 trials were no-go trials. Trial order was randomized, with the constraint that the same cue-image pair did not appear on consecutive trials.

Before starting the experiment participants performed a short practice block with trial-wise feedback to ensure good task performance. The practice block was repeated if participants failed understand the task or had poor response accuracy (< 70%) or speed (> 1000 ms). Response feedback was only provided during the practice block, however during main blocks average performance was displayed at the end of a block, both in terms of response speed and accuracy. Then participants performed six behavioral blocks before EEG recording, identical to the main task but with a shorter intertrial interval (average 2000 ms) and adjusted transitional probabilities to enhance statistical learning. Specifically, during behavioral blocks each expected pair was shown 21 times (i.e., 168 expected trials total). A short break of ~15min was inserted after three behavioral blocks to allow participants to recover and for preparations of the EEG setup. Subsequently, the EEG cap was fitted, and conductive gel applied while participants performed the remaining three behavioral blocks. Finally, participants completed eight experimental blocks during which EEG was recorded (see Figure 1C for an overview of the procedure).

### Deep Neural Network

We used an AlexNet [56] implementation pretrained on the ecoset dataset [55], to quantify visual surprise at different feature levels. For more details on the DNN we refer to Mehrer et al. [55]. Representational dissimilarity matrices (RDMs) were computed using Pearson correlation distance between the activations elicited by each image in the network layers. Layer 2 activations represented low-level visual features (e.g., edges, textures), while layer 8 activations captured high-level visual features (e.g., texture-like features). For DNN feature visualization using the present stimulus set see Richter et al. [7]. A high-resolution visualization of the RDMs, as well as the GloVe RDM (*Word Embeddings*) are shared online, see section: *Code and Data Availability*. To minimize effects due to the random initialization of network weights before training [57], we averaged RDMs of ten trained instances of the DNN with different seeds. For each unexpected trial, we then calculated the representational dissimilarity between the unexpected seen image and the expected image for the DNN layers of interest.

### Word Embeddings

To construct non-visual semantic distances, we used word embeddings derived from a pre-trained GloVe model (glove-wiki-gigaword-300; trained on: Wikipedia 2014 + Gigaword 5; [43]). For each category label (e.g. “airplane”), we added the top four semantically related terms to form a cluster of five related words. For example, a cluster was “airplane, airplanes, plane, aircraft, planes”. Labels of other image categories were excluded from the semantic cluster of the current label. For example, “car” and “airplane” could not appear in the same cluster, because both were image categories. This ensured maximal differentiation of clusters. Representations of categories were thus not limited to a single embedding but derived from the average pairwise distances between all words in their respective clusters. Pairwise distances were computed as 1 – cosine similarity between corresponding word embeddings. Distances between two different categories were calculated by averaging these pairwise distances over all word pairs drawn from the two clusters. Similarly, distances of a category to itself were determined by averaging the pairwise distances between different words within its own cluster, thereby avoiding trivial zero-distance result that arise when using a single word to represent a category.

### EEG Data Acquisition and Preprocessing

EEG data were recorded using a 64-channel actiCap Slim system (BrainVision) mounted on an elastic cap, following the international 10–20 system. Electrode impedances were lowered below 10 kΩ. EEG signals were referenced online to the FCz electrode and digitized at a sampling rate of 1,000 Hz. Electro-oculogram (EOG) was recorded with two bipolar pairs. Vertical EOG electrodes were placed above and below the eye, and horizontal EOG electrodes at the left and right outer canthi. These channels were used to detect blinks and eye movements for later artifact rejection. EEG data preprocessing was conducted using the FieldTrip toolbox [58] in MATLAB (The MathWorks, Inc., Natick, Massachusetts, United States). The preprocessing pipeline comprised the following steps:

#### Channel Inspection and Removal

Raw EEG data were visually inspected to identify and remove bad channels exhibiting excessive noise or artifacts. On average, 1.6 channels (range 0-6) were removed per participant.

#### Resampling

Continuous EEG data were down-sampled offline to 256 Hz.

#### Filtering

Continuous EEG data were notch filtered (49 to 51 Hz and 99 to 101 Hz) to remove power line noise, and band-pass filtered between 0.1 Hz and 126 Hz using a finite impulse response windowed sinc (FIRws) filter. This filter choice minimizes phase distortions and effectively attenuates frequencies outside the desired range.

#### Epoching

Data were segmented into epochs ranging from −0.8 seconds to 1 second relative to image onset, thus −0.5 seconds corresponds to cue onset. This window captured both pre-trial baseline activity and post-stimulus neural responses.

#### Baseline Correction

Baseline correction was applied using the interval from −0.7 seconds to −0.5 seconds before stimulus onset (equivalent to −0.2 seconds to 0 seconds before cue onset). This pre-cue baseline period was chosen to avoid contamination from activity following cue onset, related to stimulus anticipation and cue processing.

#### Independent Component Analysis (ICA)

ICA was performed to identify and remove components associated with physiological artifacts. The extended Infomax algorithm was used for ICA decomposition using FieldTrip [58]. Components reflecting eye blinks, lateral eye movements, muscle activity, and artifacts were identified based on their characteristic scalp topography, power spectrum, and time course. These components were removed from the data. On average, 1.7 components (range 1-3) were rejected per participant.

#### Artifact Rejection

Automated artifact rejection, as implemented in FieldTrip [58], was used to exclude trials contaminated by residual artifacts not captured by ICA. This included trials with excessive EOG activity (z score >4) or muscle artifacts (z score >8), identified using threshold-based criteria on voltage fluctuations and signal variance. An average of 797 trials (SD = 69; 77.8%) out of the original 1024 trials were retained per participant after artifact rejection.

#### Interpolation of Bad Channels

Removed channels were reconstructed using spherical spline interpolation.

#### Re-referencing

The EEG data were re-referenced to the average reference to minimize the influence of any single electrode.

#### Analysis-Specific Preprocessing

For analyses examining the univariate scaling of neural responses by visual surprise, additional preprocessing steps were applied to the epoched data, consisting of a band-pass filter between 1 Hz and 60 Hz using a FIR filter (with padding). This filtering focused on frequencies most relevant for event-related potentials (ERPs) and reduced slow drifts that could confound ERP analyses. For both univariate and multivariate analyses (RSA) a baseline correction was performed before analysis, using the 200 ms interval before cue onset (i.e. −700 ms to −500 ms relative to image onset).

### EEG Data Analysis

#### ERP Scaling Analysis

To assess the influence of surprise on visual ERPs we performed multiple linear regression analyses. In brief, for each participant and time point of the EEG signal we regressed the EEG amplitude onto the model derived dissimilarity measures (also see Figure 1D, and sections *Deep Neural Network* and *Word Embedding*). The resulting coefficients (betas) indicate how much the EEG voltage is up (positive betas) or downregulated (negative betas) as a function of surprise. The primary surprise metrics of interest were low-level visual surprise (DNN layer 2) and high-level visual surprise (DNN layer 8). We also included a non-visual semantic surprise regressor based on word embeddings (GloVe), a task regressor reflecting animacy category, and an untrained (randomly initialized) DNN layer 8 instance to account for potential biases due to network architecture. Only trials with unexpected stimuli and correct responses (hits) were included in this analysis. Because our focus was on visually evoked ERPs we averaged the obtained beta coefficients across occipital and parieto-occipital electrodes (Oz, O1, O2, POz, PO7, PO8, PO3, PO4). We chose to average across all available occipital and parieto-occipital electrodes to increase the robustness of our results and to avoid concerns arising from selectively focusing analyses on individual electrodes. However, we did restrict our analysis to parieto-occipital electrodes given our focus on visual processing in the present study [12,15–21,23–25]. The beta coefficients were then smoothed for each participant using a Savitzky–Golay filter (order = 2; window length = 9 samples, corresponding to ~35 ms) before being submitted to group level inference. Cluster-based permutation tests (100,000 permutations), as implemented in FieldTrip [58], were conducted for each model (layer 2, layer 8, GloVe, Animacy, untrained layer 8) contrasting the obtained beta coefficients against zero (no modulation).

#### Individual ERP Peak Analysis

In addition to the scaling analysis outlined above, we performed a variant of the approach allowing for individual variations of the ERP peak timing. In this analysis we determined for each participant five key visually evoked ERPs (P1, N1, P2, N2, P3). These ERPs were defined on each participant’s averaged ERP amplitude over parieto-occipital sensors as the maximal positive (P1, P2, P3) or negative (N1, N2) individual deflection within time windows commonly reported in the literature for each ERP. We used the following post-stimulus onset time windows for the ERPs: P1: 80-130ms; N1/N170: 130-190ms; P2: 180-250ms; N2: 200-300ms, P3: 300-600ms post-stimulus onset [12–25]. Once the maximal individual deflections were determined we took a 20ms window around the peak and averaged the individual surprise beta coefficients (see: *ERP Scaling Analysis*) within the time window. Thus, the resulting beta coefficients reflect the up or down regulation of visually evoked responses by the five surprise models at the individual ERP peaks. We then performed group level inference on these individual betas and computed the mean across participants. Resulting p values of one-sample t-tests were FDR corrected for total number of tests (ERPs x Models), while paired t-tests were FDR corrected for the number of models, since no paired tests were performed between ERPs.

#### Representational Similarity Analysis (RSA)

Before performing the key analyses outlined above, we first ensured that the model derived representations reliably explained variance in the EEG signal, independent of expectations. To this end, we performed a RSA [10]. First we created the model derived RDMs by computing all pair-wise distances (1 – correlation) between stimuli using AlexNet (layer 2 and layer 8), GloVe, animacy category (0 or 1), and a randomly initialized instance of AlexNet (also see Figure 1D, *Deep Neural Network* and *Word Embedding*). For each participant this results in an 8 by 8 RDM for each model of interest, representing the pairwise representational distances between the 8 specific stimuli seen by that participant. Next, for each participant we computed the neural RDMs per timepoint of the EEG by calculating the pair-wise distances (1 – Pearson correlation) between stimuli across all sensors and EEG blocks. Unlike in the primary analysis (see: *ERP Scaling Analysis*) this analysis included all trials, irrespective of expectation status. We then correlated each of the five model RDMs and the neural RDMs per timepoint and participant using Kendall’s Tau. Therefore, the resulting timeseries of correlation coefficient indicates for each model the shared representation geometry of the neural signal and the model RDM. Finally, after smoothing using a Savitzky–Golay filter (order = 2; window length = 9 samples, corresponding to ~35 ms), we subjected the correlation coefficients to group level inference using cluster-based permutation tests, contrasting the correlation against zero (no shared representational geometry).

#### RSA Time Analysis

Using the correlation coefficients per participant from the RSA outlined above we additionally estimated the onset time of the modulations. This analysis tests the expectation that low-level visual features ought to modulate visually evoked responses before higher-level features. In brief, we used a jackknife procedure that restores individual estimates [59] to obtain individual onset latencies (50% of maximum amplitude) within the first 200ms post stimulus onset using the latency toolbox [60]. We computed group averages and performed group level statistics by subjecting the transformed jackknife estimates [59] to paired t-tests. A similar approach was used to estimate the onset latency of the ERP scaling analysis using beta coefficients per surprise regressor.

### Behavioral Data Analysis

RTs and accuracy were calculated for expected and unexpected trials, with the latter split into trials requiring the same or different response as the expected stimulus. Trials with RTs shorter than 100 ms or longer than 1500 ms were excluded. Only trials with correct responses (hits) were included in RT analyses. Behavioral data were submitted to repeated measures ANOVAs. Posthoc paired t-tests compared RTs and accuracy between conditions. P values are corrected for multiple comparisons using the Holm– Bonferroni method as implemented in JASP [61].

### Statistical Analysis and Software

Statistical analyses were performed using MATLAB (The MathWorks, Inc., Natick, Massachusetts, United States) and the FieldTrip toolbox [58]. For ERP data, cluster-based permutation tests were employed to control for multiple comparisons across time points and electrodes. Beta coefficients from the individual peak analysis were tested against zero using one-sample t-tests. Additionally, for this analysis we used equivalent Bayesian statistical tests to evaluate not statistically significant results, computing Bayes Factors to assess evidence for the absence of effects. Bayesian tests were performed using JASP 0.19.3.0 with default priors; i.e. using a Cauchy prior of 0.707 for Bayesian one-sample t-tests. Qualitative interpretation of Bayes Factors were based on [62].

## Code and Data Availability

All data and code will be made available upon publication in a peer-reviewed journal. Supplemental visualization of RDMs is available here: https://doi.org/10.17605/OSF.IO/QSRZ4.

## Ethics Statement

All participants provided written informed consent before participating, compensated at a rate of €10 per hour, and fully debriefed after completion of the experiment. The study followed guidelines set by the Declaration of Helsinki, as well as institutional guidelines, and the was approved by the local ethics committee (reference 3270/CEIH/2023).

## Acknowledgements

We thank Tijana Baclic for assistance with data acquisition.

## Contributions

*David Richter*: Conceptualization, Data curation, Formal analysis, Funding acquisition, Investigation, Methodology, Project administration, Resources, Software, Supervision, Validation, Visualization, Writing – original draft, Writing – review & editing.

*Paula Pena*: Data curation, Investigation, Methodology, Writing – review & editing.

*María Ruz*: Conceptualization, Funding acquisition, Project administration, Resources, Supervision, Writing – review & editing.

## Supporting Information

**S1 Figure.**
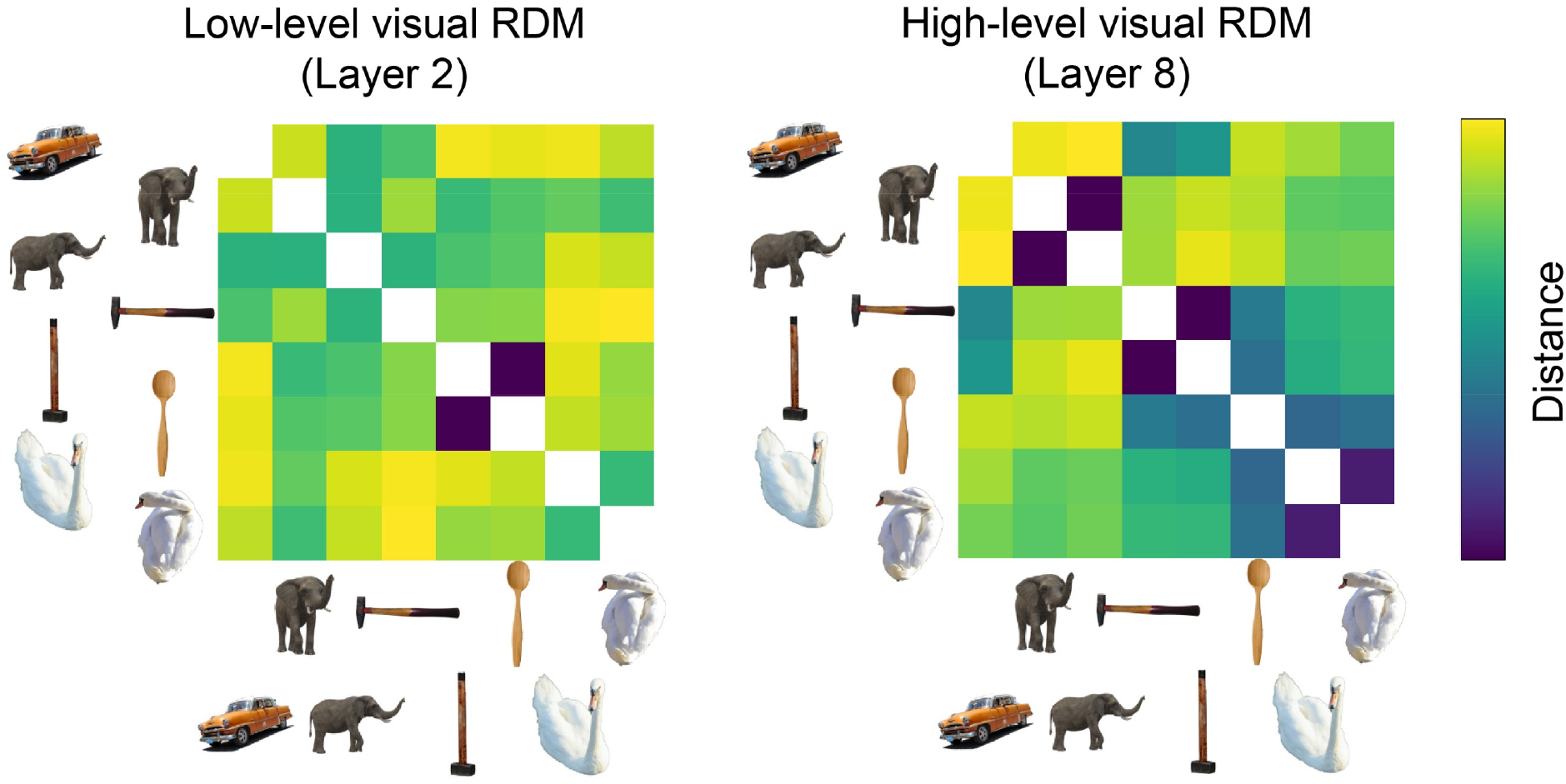
Stimulus set and RDM visualization of one example participant. Depicted is the stimulus set, comprised of eight images, and corresponding RDMs from DNN layer 2 (left) and layer 8 (right) for one example participant. The example demonstrates several aspects of stimulus selection. Frist, each participant saw four animate and four inanimate stimuli. Second, the stimulus selection emphasizes large variance within RDMs and low correlation between the two RDMs. This resulted in stimulus sets of large low-level visual variability (e.g. objects of different orientations, shapes, and colours) and high-level variability (e.g. different types of objects and animals). Third, the stimulus selection exploits systematic changes in distance between RDMs. For example, in the case depicted here, we see that the two elephants were distant in low-level visual space, yet close in high-level space. In contrast, the vertical hammer and spoon were close in low-level visual space, but more distant in high-level space. Combined this selection ensures significant variability in both low-level and high-level visual features, approximating the fact that we encounter diverse objects during natural object perception, while simultaneously optimizing stimulus selection for the planned analyses. However, we stress that the stimuli and RDMs depicted here only constitute one example. Importantly, our results are not contingent on this specific stimulus set or RDMs, because each participant saw a different set of images and cue-stimulus associations, selected from an image database of 233 stimuli (for details see section *Stimuli and Experimental Paradigm*). Image sets and RDM figures of the remaining participants are available online: https://doi.org/10.17605/OSF.IO/QSRZ4.

**S2 Figure.**
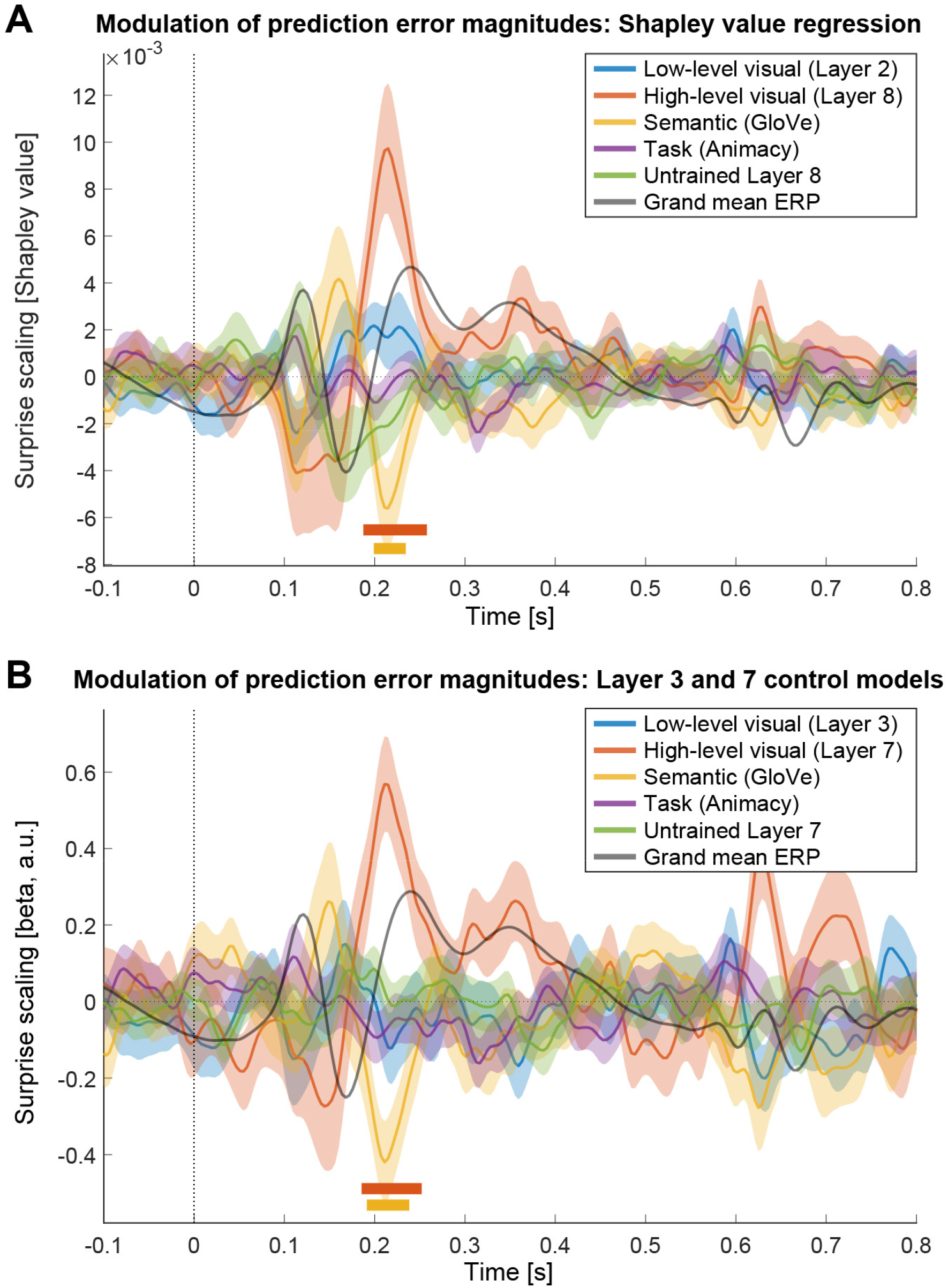
Control analyses. **A)** Control analysis using Shapley regression. To address potential concerns arising from correlation between predictors, we repeated the surprise scaling analysis using Shapley value regression. In brief, it computes the average marginal contribution over all subsets of models for the five predictors, thereby helping to address possible issues arising from multicollinearity. While Shapley values are unsigned, we reconstructed signs from the original coefficients. Results replicated those reported in our primary analysis using ordinary least squares regression, showing a pronounced positive modulation of neural responses by high-level visual surprise and a slight negative modulation by semantic surprise. Low-level visual surprise again did not scale responses. **B)** Control analysis using neighboring layers of the primary surprise scaling analysis. To ensure that results were not contingent on the (a priori) selected DNN layers, we repeated the surprise scaling analysis using neighboring layers. Thus, in this analysis layer 3 was used as low-level and layer 7 as high-level visual surprise model. Note, in our implementation of AlexNet layer 8 is the final layer before softmax, thus the neighboring layer for layer 8 containing high-level visual representations is layer 7. Results replicated the primary findings, showing a clear upregulation of visually evoked responses by high-level visual surprise approximately 200 ms after stimulus onset. As in the main analysis, we again found no modulation of ERP amplitudes by low-level visual surprise (here layer 3). Therefore, this control analysis demonstrates that our results are not contingent on the precise layers selected for analysis but replicate across neighboring layers, reinforcing our conclusion that visually evoked responses are primarily modulated by high-level visual surprise. Colored bars above the abscissa denote statistically significant clusters (*p*_cluster_ < 0.05).

**S1 Table.** Results of paired t-tests contrasting the modulation ERP magnitudes by high-level visual surprise (layer 8) against the four other surprise models. ERP time windows were defined per participant using the maximal positive deflection within commonly reported time windows for the P2 potential. Beta coefficients were averaged per participant within these time windows and contrasted between models on the group level. Test statistics reflect paired t-tests. P values are FDR corrected using the Benjamini and Hochberg method. Effect sizes are given as Cohen’s *d*_z_.

| ERP | Contrast | Test statistic: $t_{(37)}$ | P value | Effect size: $d_z$ |
| --- | --- | --- | --- | --- |
| P2 | Layer 8 vs Layer 2 | 3.79 | 0.002 | 0.61 |
| P2 | Layer 8 vs GloVe | 5.56 | 2.5e-6 | 0.90 |
| P2 | Layer 8 vs Animacy | 3.72 | 0.001 | 0.60 |
| P2 | Layer 8 vs Control model | 5.43 | 3.8e-6 | 0.88 |

**S2 Table.** Results of one-sample t-tests contrasting the modulation of ERP magnitudes by surprise against zero (no modulation). ERP time windows were defined per participant using the maximal negative (N1, N2) or positive (P1, P2, P3) deflection within commonly reported time windows of the respective ERP. Beta coefficients were averaged per participant within these time windows and contrasted against zero at the group level. Test statistics reflect one-sample t-tests. P values are FDR corrected using the Benjamini and Hochberg method. Effect sizes are given as Cohen’s *d*_z_. Bayes factors are reported as BF10. *p*_corrected_ < 0.05 are marked in bold.

| ERP | Surprise model | Test stat.: $t_{(37)}$ | P value | Effect size: $d_z$ | BF<sub>10</sub> |
| --- | --- | --- | --- | --- | --- |
| P1 | Low-level visual (layer 2) | -2.47 | 0.077 | -0.40 | 2.473 |
| P1 | High-level visual (layer 8) | -0.63 | 0.703 | -0.10 | 0.210 |
| P1 | Semantic (GloVe) | -0.76 | 0.669 | -0.12 | 0.228 |
| P1 | Task (Animacy) | 0.14 | 0.893 | 0.02 | 0.176 |
| P1 | Control (untrained layer 8) | 0.65 | 0.703 | 0.10 | 0.212 |
| N1 | Low-level visual (layer 2) | 1.13 | 0.478 | 0.18 | 0.314 |
| N1 | High-level visual (layer 8) | -1.57 | 0.314 | -0.25 | 0.535 |
| N1 | Semantic (GloVe) | 2.01 | 0.183 | 0.33 | 1.071 |
| N1 | Task (Animacy) | -0.35 | 0.823 | -0.06 | 0.185 |
| N1 | Control (untrained layer 8) | -2.89 | 0.054 | -0.47 | 6.019 |
| P2 | Low-level visual (layer 2) | 0.31 | 0.823 | 0.05 | 0.183 |
| **P2** | **High-level visual (layer 8)** | **5.46** | **8.4e-5** | **0.89** | **5578.153** |
| P2 | Semantic (GloVe) | -2.85 | 0.054 | -0.46 | 5.605 |
| P2 | Task (Animacy) | -0.89 | 0.617 | -0.14 | 0.253 |
| P2 | Control (untrained layer 8) | -1.50 | 0.322 | -0.24 | 0.489 |
| N2 | Low-level visual (layer 2) | 1.37 | 0.373 | 0.22 | 0.413 |
| N2 | High-level visual (layer 8) | 2.68 | 0.054 | 0.44 | 3.883 |
| N2 | Semantic (GloVe) | -1.94 | 0.189 | -0.31 | 0.938 |
| N2 | Task (Animacy) | 0.26 | 0.827 | 0.04 | 0.180 |
| N2 | Control (untrained layer 8) | -1.22 | 0.446 | -0.20 | 0.345 |
| P3 | Low-level visual (layer 2) | 0.39 | 0.823 | 0.06 | 0.188 |
| P3 | High-level visual (layer 8) | 2.77 | 0.054 | 0.45 | 4.692 |
| P3 | Semantic (GloVe) | -1.70 | 0.273 | -0.28 | 0.643 |
| P3 | Task (Animacy) | -0.86 | 0.617 | -0.14 | 0.246 |
| P3 | Control (untrained layer 8) | 0.38 | 0.823 | 0.06 | 0.187 |

